# Comparative Chloroplast Genomics of Henbit (*Lamium amplexicaule*) Reveals Diagnostic Plastid Markers in *Lamium*

**DOI:** 10.64898/2026.11.25.769096

**Authors:** Ryan Wheeler, Ema de Alba, Lillie Hanson, Lexi Avila, Gerardo Laureano, Christina Johne, Alok Arun

## Abstract

*Lamium amplexicaule* (henbit deadnettle) is a widespread member of Lamiaceae, but plastome-level comparisons within *Lamium* are still limited. In this study, we characterized the complete chloroplast genome of an *L. amplexicaule* accession collected in Turlock, California, and compared it with other available *Lamium* plastomes. The *L. amplexicaule*_Turlock plastome was 150,642 bp in length and had the typical quadripartite structure, with 113 distinct chloroplast genes. Phylogenomic analysis based on 79 shared protein-coding genes across 31 taxa placed the three sampled *L. amplexicaule* accessions together in a strongly supported clade. Comparative analysis of 13 *Lamium* plastomes showed that gene content, IR structure, and overall genome organization were highly conserved. The three *L. amplexicaule* plastomes differed by 36 SNPs and 16 inferred indel events. Codon usage was biased toward A- and U-ending codons, and Ka/Ks analysis showed mainly purifying selection, with no significant evidence of positive selection. Sliding-window analysis identified *ycf1* and the *petA-psbJ* region as the most variable regions in the dataset. Both regions showed good discrimination among the seven sampled *Lamium* species in the *in-silico* analysis. These results provide a new chloroplast genome for *L. amplexicaule* and show that *Lamium* plastomes are highly conserved overall but still contain useful sequence variation for phylogenetic and taxonomic studies.

## 1. Introduction

Chloroplast genomes are widely used in plant phylogenetic and comparative studies because they are relatively small, conserved, and contain useful genetic information. Most angiosperm chloroplast genomes have a typical quadripartite structure consisting of a large single-copy (LSC) region, a small single-copy (SSC) region, and two inverted repeat regions (IRa and IRb) [1]. Although chloroplast genomes are generally conserved, differences can occur in gene content, repeat sequences, IR boundaries, and nucleotide sequence. Complete plastome data can therefore be useful for studying both phylogenetic relationships and genome variation.

Lamiaceae is a large and diverse plant family, and chloroplast genomes have been used to study relationships among its major groups. Zhao et al. [2] used 79 plastid protein-coding genes to examine phylogenetic relationships across the family. Within subfamily Lamioideae, the genus *Lamium* includes annual and perennial species and has a complex evolutionary history [3,4]. Previous work has found disagreement between chloroplast and nuclear phylogenies and evidence of hybridization and polyploidy within the genus [4]. *Lamium amplexicaule* is an annual species that produces both chasmogamous and cleistogamous flowers, and the proportion of these flower types can change under different environmental conditions [5].

Complete chloroplast genomes have been reported for several *Lamium* species, including *L. takesimense*, *L. amplexicaule*, and *L. barbatum* [2,6,7]. However, comparisons involving several *L. amplexicaule* accessions and a larger group of *Lamium* species are still limited. Earlier work also did not recover all sampled *L. amplexicaule* material as a single lineage [4], making additional plastome sampling useful for examining its phylogenetic placement.

Chloroplast genomes can also be used to identify regions that vary among closely related taxa. Simple sequence repeats are commonly used as markers in plant genetic studies [8], and several chloroplast regions, including *psbJ-petA*, *rps16-trnQ*, *psbE-petL*, *ndhF*, *rpoB-trnC*, and *ycf1*, have been reported as variable at low taxonomic levels [9,10]. The *ycf1* gene has also shown strong potential as a plastid barcode region [11]. Other comparisons, including IR boundaries, codon usage, SNPs and indels, and Ka/Ks ratios, can provide additional information on plastome variation and evolution.

In this study, we characterized the complete chloroplast genome of *L. amplexicaule* collected in Turlock, California. Additionally, a total of 31 plastomes were used for phylogenomic analysis, including 13 *Lamium* accessions from NCBI database. The *Lamium* plastomes were compared for gene content, synteny, repeat sequences, codon usage, IR boundaries, nucleotide diversity, SNPs and indels, and selection pressure. We also examined gene-tree concordance, tested the effect of removing *clpP1* from the phylogenomic dataset, and evaluated variable regions as possible diagnostic markers.

## 2. Materials and methods

### 2.1 Plant material, DNA extraction, sequencing, assembly, and annotation

Specimens of *Lamium amplexicaule* were collected on the California State University Stanislaus campus in Turlock, California, USA, at approximately 37°31′32.0″N, 120°51′22.7″W in February 2026. Whole plants were collected and separated into stems, leaves, roots, immature flowers, and mature flowers. Genomic DNA was extracted from young leaf tissue using a Qiagen DNA extraction kit following manufacturer’s protocol. DNA quality and concentration were assessed prior to sequencing. The extracted DNA was used to prepare sequencing libraries, which were sequenced on an Illumina NovaSeq 6000 platform to generate high-throughput sequencing data. The sequencing yielded a total of 210,033,128 reads. Raw reads were quality filtered using fastp v1.3.3 [14]. The chloroplast genome was assembled using GetOrganelle v1.7.7.1 with the embplant_pt database and k-mer sizes of 21, 45, 65, 85, 105, and 115 [15]. The completed plastome was annotated using GeSeq with the Plastid (land plants) profile and Chloë annotation support [12]. The raw reads were deposited to NCBI BioProject PRJNA1564337 and can be accessed at Figshare link: 10.6084/m9.figshare.35063311

### 2.2 Taxon sampling and comparative plastome dataset

*L. amplexicaule* plastome was compared with complete chloroplast genomes obtained from NCBI GenBank. The final dataset contained 31 plastomes, including 13 *Lamium* accessions, 10 additional members of Lamioideae, six representatives of other Lamiaceae lineages, and two external Lamiales outgroups.

The *Lamium* comparative dataset included *L. amplexicaule* collected from Turlock, *L. amplexicaule* MT473770, *L. amplexicaule* PQ231089, and representatives of *L. album*, *L. album* subsp. *album*, *L. barbatum*, *L. purpureum*, *L. purpureum* var. *purpureum*, *L. takesimense*, *L. galeobdolon*, *L. galeobdolon* subsp. *flavidum*, and *L. orvala*. *Mazus pumilus* MT937187 and *Phryma leptostachya* subsp. *asiatica* MT948145 were used as external outgroups. The complete taxon list and GenBank accession numbers are provided in (S1_Table).

All 31 complete chloroplast genomes included in the dataset were reannotated using a standardized approach and incorporated into the phylogenomic analysis to examine relationships among the sampled taxa. Comparative analyses of genome structure, gene content, repeat sequences, codon usage, inverted-repeat boundaries, nucleotide variation, SNPs, indels, and selection pressure were conducted using the 13 plastomes representing species within the genus *Lamium*. Supporting numerical data for these analyses are provided in (S1_Data).

### 2.3 Standardized plastome annotation and annotation quality assessment

All 31 plastomes were reannotated using GeSeq with the Plastid (land plants) profile and Chloë annotation support [12]. Protein-coding genes, tRNAs, rRNAs, intron-containing genes, trans-spliced *rps12*, partial gene fragments, and IR features were checked.

Duplicated IR copies were collapsed for gene-count comparisons, and partial boundary fragments were not counted as additional genes. Only Chloë-supported tRNA annotations were retained. ARAGORN-only tRNA predictions were excluded. The same gene names and counting method were used for all accessions.

### 2.4 Phylogenomic analysis, gene-tree concordance, and *clpP1* sensitivity analysis

Protein-coding sequences were extracted using PhyloSuite [16]. Duplicate IR copies and partial gene fragments were removed, leaving 79 protein-coding genes shared among all 31 taxa. Each gene was aligned separately using MACSE with Genetic Code Table 11 and default settings [17]. The alignments were trimmed using trimAl with the automated1 method [18] and concatenated in PhyloSuite. The final nucleotide matrix was 68,112 bp long.

Maximum-likelihood phylogenetic analysis was performed using IQ-TREE v3.0.1 [19]. ModelFinder was used for substitution-model selection and partition optimization [20]. An edge-linked proportional partition model was used [21]. Branch support was calculated using 5,000 ultrafast bootstrap replicates and 1,000 SH-like approximate likelihood-ratio test replicates [22,23]. The tree was rooted using *M. pumilus* and *P. leptostachya* subsp. *asiatica*. Separate maximum-likelihood trees were also generated for each of the 79 protein-coding genes. Gene concordance factors were calculated in IQ-TREE using the concatenated tree as the reference [13].

The *clpP1* gene contained a localized coding-frame/annotation concern. To test whether this gene affected the phylogenetic result, the 591-bp *clpP1* partition was removed and the analysis was repeated. The reduced matrix contained 78 genes and 67,521 aligned positions. The same IQ-TREE analysis and branch-support methods were used, and gene concordance factors were recalculated from the remaining 78 gene trees.

### 2.5 Whole-plastome synteny analysis

Whole-plastome synteny was compared among six *Lamium* plastomes using AliTV v1.0.6 [24]. Pairwise genome alignments were produced with LASTZ v1.04.52 [25]. The analysis included the *L. amplexicaule*_Turlock plastome, *L. amplexicaule* MT473770, *L. album* OR565909, *L. barbatum* PV802395, *L. purpureum* var. *purpureum* PZ055112, and *L. takesimense* MN240520. Alignments were generated in both sequence orientations. Homologous regions with 85–100% sequence identity and a minimum alignment length of 1,061 bp were displayed. Only links between adjacent plastomes were shown. Collinear and reverse-orientation regions were compared, with particular attention to the IR/SSC region.

### 2.6 Comparative gene-content analysis

Gene content was compared among the 13 *Lamium* plastomes using the standardized GeSeq annotations. Protein-coding genes, tRNAs, and rRNAs were recorded for each accession. Duplicated IR copies were collapsed, and partial boundary annotations were grouped with their corresponding full-length genes. Only Chloë-supported tRNAs were retained. Genes were grouped into photosynthesis, self-replication, other conserved plastid functions, and genes of uncertain function.

### 2.7 Repeat sequence analyses

Simple sequence repeats were identified using MISA-web [26]. The minimum repeat numbers were 10 for mononucleotide repeats, five for dinucleotide repeats, four for trinucleotide repeats, and three for tetra-, penta-, and hexanucleotide repeats. The maximum distance between SSRs classified as a compound repeat was 100 bp.

Long dispersed repeats were identified using Vmatch [27]. A minimum repeat length of 30 bp and a Hamming distance of 3 were used. Matches representing the normal plastome inverted repeats were removed before repeat totals were calculated. Overlapping matches from the same repeat locus were collapsed, and the longest match was retained.

Tandem repeats were identified using Tandem Repeats Finder v4.09 [28]. Parameters were set to a match score of 2, mismatch penalty of 7, indel penalty of 7, match probability of 80, indel probability of 10, minimum alignment score of 50, and maximum period size of 500 bp. Overlapping calls from the same repeat array were collapsed into one locus.

### 2.8 Codon-usage analysis

Codon usage was analyzed among the 13 *Lamium* plastomes using CodonW v1.4.2 [29]. A total of 78 shared protein-coding genes were used for the analysis. *clpP1* was excluded because of the coding-frame/annotation concern. Terminal stop codons were removed before the genes were concatenated. Relative synonymous codon usage, effective number of codons, overall GC content, GC3s, and GC content at the first, second, and third codon positions were calculated. Codons with mean RSCU values greater than 1.0 were considered preferentially used.

### 2.9 Inverted-repeat boundary analysis

IR boundaries were compared among the 13 *Lamium* plastomes using IRscope [30]. Plastome orientation and region assignments were checked before the analysis. Circular sequence origins were rotated where needed, and SSC orientation was normalized for comparison. The LSC/IRb, IRb/SSC, SSC/IRa, and IRa/LSC junctions were examined. The positions of *rps19*, *rpl2*, *ndhF*, *ycf1*, and *trnH-GUG* near the junctions were also compared.

### 2.10 Sliding-window nucleotide-diversity analysis

The 13 complete *Lamium* plastomes were aligned using MAFFT in PhyloSuite [16,31]. Plastome orientation was standardized before alignment. Nucleotide diversity was calculated using DnaSP v6.12.03 [32] with a window length of 600 bp and a step size of 200 bp. Sites containing gaps or missing data were excluded from individual window calculations. Nucleotide-diversity values were plotted across the alignment. Regions containing local diversity peaks were assigned to genes or intergenic regions using the *L. amplexicaule*_Turlock annotation and were used for later candidate-marker analysis.

### 2.11 SNP and indel analysis

Single Nucleotide Polymorphism (SNP) and Insertion Deletion (indel) variation was examined using the whole-plastome MAFFT alignment. Analyses were performed across all 13 *Lamium* plastomes and separately among the three *L. amplexicaule* accessions. SNPs were identified using SNP-sites v2.5.1 [33]. For the 13-taxon comparison, sites containing gaps or ambiguous bases were excluded. Indels were analyzed using DnaSP v6.12.03 under Model 4 (Multiallelic) [32]. For the three *L. amplexicaule* plastomes, inferred indel events were also mapped to physical gap positions in the alignment.

SNP and indel coordinates were converted to *L. amplexicaule*_Turlock plastome coordinates and classified as coding, intronic, or intergenic. Coding SNPs were classified as synonymous or nonsynonymous. Coordinates in the SSC region were converted back to the deposited *L. amplexicaule*_Turlock orientation before annotation.

### 2.12 Ka/Ks selection analysis

Ka/Ks analysis was performed using KaKs_Calculator v3.0 and the modified Yang-Nielsen method [34,35]. The same 78 shared protein-coding genes used for the codon-usage analysis were included, with *clpP1* excluded. All pairwise comparisons among the 13 accessions were analyzed for each gene, giving 6,084 gene-by-pair comparisons. Ka, Ks, and Ka/Ks were calculated for each comparison. Comparisons without a valid Ka/Ks value were treated as missing. Ka/Ks < 1 with Fisher *P* < 0.05 was considered evidence of purifying selection, while Ka/Ks > 1 with *P* < 0.05 was required for evidence of positive selection.

### 2.13 *In silico* barcode analysis

Eight 600-bp regions centered on nucleotide-diversity peaks were evaluated as candidate plastid markers. A custom Python script was used to calculate variable sites, parsimony-informative sites, haplotypes, interspecific sequence sharing, and uncorrected pairwise p-distances from complete-case sites. Following established plant DNA-barcoding approaches, barcode gaps were evaluated by comparing minimum interspecific and maximum intraspecific p-distances [43]. Barcode gap was calculated as the minimum interspecific p-distance minus the maximum intraspecific p-distance (S1_Code).

The two highest-performing regions, *ycf1* and *petA-psbJ*, were used for degenerate primer design. Primer binding was screened across the 13 orientation-normalized plastomes, and binding sites and predicted amplicon lengths were recorded. Predicted amplicons were compared for sequence variation, haplotypes, interspecific sequence sharing, and barcode-gap performance. These analyses were considered in silico screening only.

## 3. Results

### 3.1 Plastome structure and conserved gene content

The complete chloroplast genome of the *L. amplexicaule*_Turlock was 150,642 bp in length and had the typical quadripartite structure (Fig 1). The large single-copy (LSC) region was 82,812 bp, the small single-copy (SSC) region was 17,470 bp, and each inverted repeat (IR) was 25,180 bp. The overall GC content was 38.53%.

**Fig 1.**
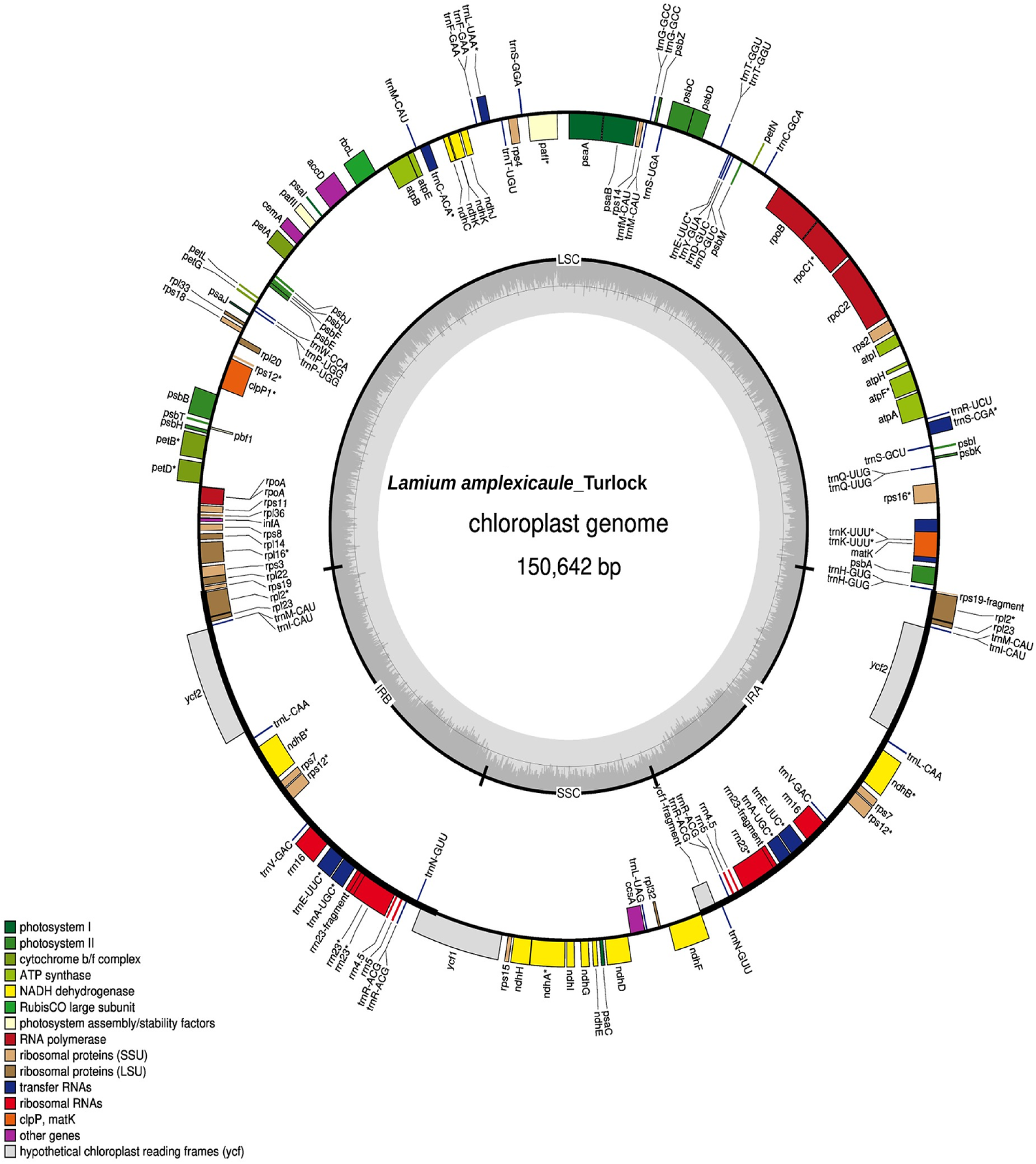
Chloroplast genome map of the *L. amplexicaule*_Turlock. The complete plastome is 150,642 bp. Genes inside and outside the circle are transcribed in opposite directions, and colors indicate functional groups. The inner circle shows GC content and the LSC, SSC, IRa, and IRb regions.

A total of 113 distinct chloroplast genes were identified after duplicated IR copies and partial gene fragments were removed. These included 79 protein-coding genes, 30 tRNA genes, and four rRNA genes. The same gene set was found in all 13 *Lamium* plastomes (Table 1). Nineteen genes contained introns, and *rps12* was trans-spliced.

**Table 1.** Chloroplast gene classification in *Lamium* plastome.

| Category | Gene group | Gene names |
|---|---|---|
| Photosynthesis | Photosystem I and assembly | psaA, psaB, psaC, psaI, psaJ, pafI<sup>a</sup>, pafII |
|  | Photosystem II and assembly | pbf1, psbA, psbB, psbC, psbD, psbE, psbF, psbH, psbI, psbJ, psbK, psbL, psbM, psbT, psbZ |
|  | NADH dehydrogenase | ndhA<sup>a</sup>, ndhB<sup>a</sup>, ndhC, ndhD, ndhE, ndhF, ndhG, ndhH, ndhI, ndhJ, ndhK |
|  | Cytochrome b6/f complex | petA, petB<sup>a</sup>, petD<sup>a</sup>, petG, petL, petN |
|  | ATP synthase | atpA, atpB, atpE, atpF<sup>a</sup>, atpH, atpI |
|  | Large subunit of Rubisco | rbcL |
| Self-replication | Large ribosomal proteins | rpl2<sup>a</sup>, rpl14, rpl16<sup>a</sup>, rpl20, rpl22, rpl23, rpl32, rpl33, rpl36 |
|  | Small ribosomal proteins | rps2, rps3, rps4, rps7, rps8, rps11, rps12<sup>ab</sup>, rps14, rps15, rps16<sup>a</sup>, rps18, rps19 |
|  | DNA-dependent RNA polymerase | rpoA, rpoB, rpoC1<sup>a</sup>, rpoC2 |
|  | Ribosomal RNAs | rrn4.5, rrn5, rrn16, rrn23<sup>a</sup> |
|  | Transfer RNAs | trnA-UGC<sup>a</sup>, trnC-GCA, trnD-GUC, trnE-UUC, trnF-GAA, trnG-GCC, trnG-UCC<sup>a</sup>, trnH-GUG, trnI-CAU, trnI-GAU<sup>a</sup>, trnK-UUU<sup>a</sup>, trnL-CAA, trnL-UAA<sup>a</sup>, trnL-UAG, trnM-CAU, trnN-GUU, trnP-UGG, trnQ-UUG, trnR-ACG, trnR-UCU, trnS-GCU, trnS-GGA, trnS-UGA, trnT-GGU, trnT-UGU, trnV-GAC, trnV-UAC<sup>a</sup>, trnW-CCA, trnY-GUA, trnfM-CAU |
| Other genes | Maturase | matK |
|  | Protease | clpP1<sup>a</sup> |
|  | Envelope membrane protein | cemA |
|  | Acetyl-CoA carboxylase subunit | accD |
|  | Cytochrome c synthesis protein | ccsA |
|  | Translation initiation factor | infA |
| Conserved open reading frames | Conserved genes | ycf1, ycf2 |

<sup>a</sup> Gene containing one or more introns; <sup>b</sup> trans-spliced gene.

### 3.2 Phylogenomic relationships and gene-tree concordance

The 79 protein-coding gene datasets contained 68,112 aligned nucleotide positions across 31 taxa (Fig 2). The sampled *Lamium* plastomes formed a strongly supported clade (SH-aLRT/UFBoot = 100/100; gCF = 59.49%). The three *L. amplexicaule* accessions also formed a strongly supported clade (100/100; gCF = 77.22%). MT473770 and PQ231089 were recovered as sister accessions (97.9/100; gCF = 37.97%), with the *L. amplexicaule*_Turlock sister to this pair.

**Fig 2.**
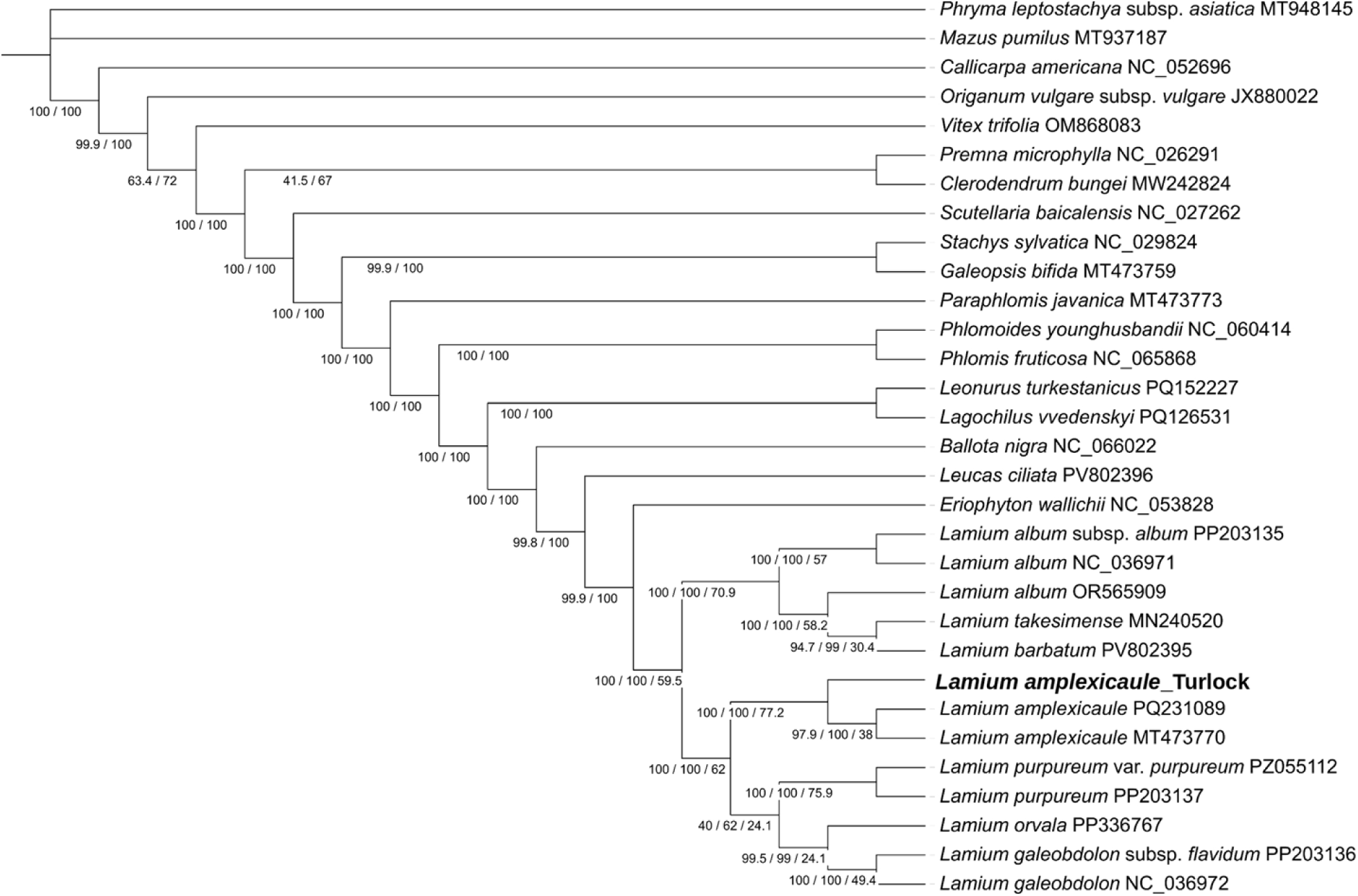
Maximum-likelihood phylogeny of *Lamium amplexicaule* and related Lamiales based on 79 chloroplast protein-coding genes. Node values show SH-aLRT/UFBoot/gCF support where available. *Mazus pumilus* and *Phryma leptostachya subsp. asiatica* were used as outgroups. The *L. amplexicaule*_Turlock accession is shown in bold.

The two *L. purpureum* accessions were recovered together (100/100; gCF = 75.95%), as were the two *L. galeobdolon* accessions (100/100; gCF = 49.37%). *Lamium barbatum* and *L. takesimense* were also recovered as sisters (94.7/99; gCF = 30.38%). One internal *Lamium* branch had lower support (40/62; gCF = 24.05%).

Removal of the 591-bp *clpP1* partition produced a 78-gene matrix containing 67,521 aligned positions. The 79-gene and 78-gene trees contained the same 28 internal bipartitions, with a Robinson-Foulds distance of 0. Gene concordance factors changed by less than one percentage point for the major branches. The *Lamium* clade changed from 59.49% to 58.97%, the *L. amplexicaule* clade from 77.22% to 76.92%, and the MT473770 + PQ231089 pair from 37.97% to 38.46%.

### 3.3 Whole-plastome synteny

The six *Lamium* plastomes showed a high level of collinearity (Fig 3). Reverse-orientation homologous regions were mainly found around the IR/SSC region, including the opposite SSC orientation of the *L. amplexicaule*_Turlock plastome. No large rearrangements were detected outside this region. The SSC orientation of the *L. amplexicaule*_Turlock plastome was therefore considered an alternative plastome structural isomer rather than a fixed inversion.

**Fig 3.**
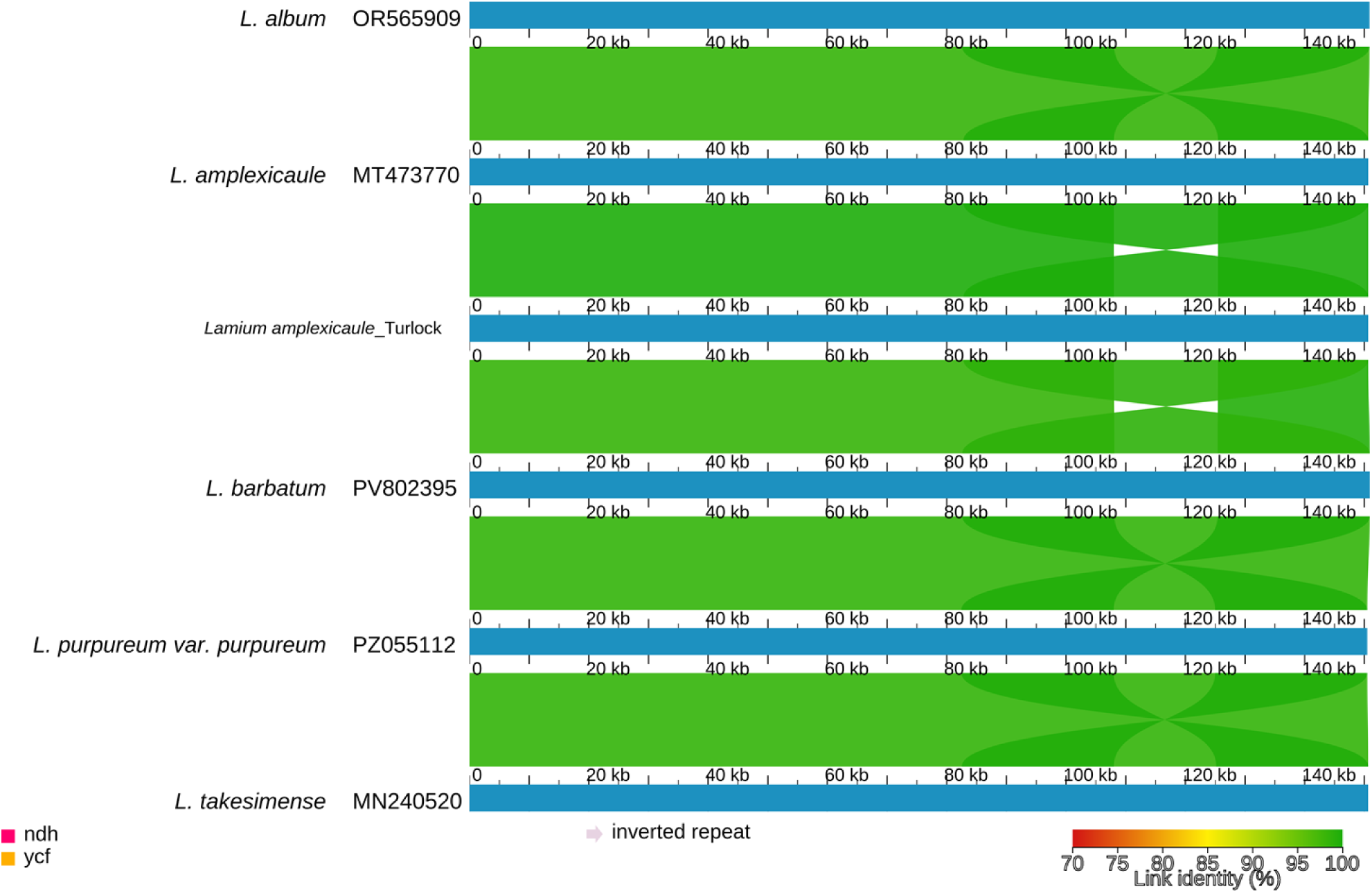
Whole-plastome synteny among six *Lamium* chloroplast genomes. Homologous regions were identified with LASTZ and visualized in AliTV. Crossed links indicate reverse-orientation homology, mainly around the IR/SSC region.

### 3.4 Inverted-repeat boundary variation

The IR boundaries were similar among the 13 *Lamium* plastomes (Fig 4). The genes *rps19*, *rpl2*, *ycf1*, *ndhF*, and *trnH* occurred in similar positions around the four junctions. IR lengths ranged from 25,180 to 25,553 bp. All three *L. amplexicaule* plastomes had IR regions of 25,180 bp, the shortest in the dataset. The *L. amplexicaule*_Turlock plastome had an 82,812-bp LSC and a 17,470-bp SSC. MT473770 and PQ231089 each had an 82,813-bp LSC and a 17,477-bp SSC. Only small differences in IR expansion and contraction were observed among the other *Lamium* plastomes.

**Fig 4.**
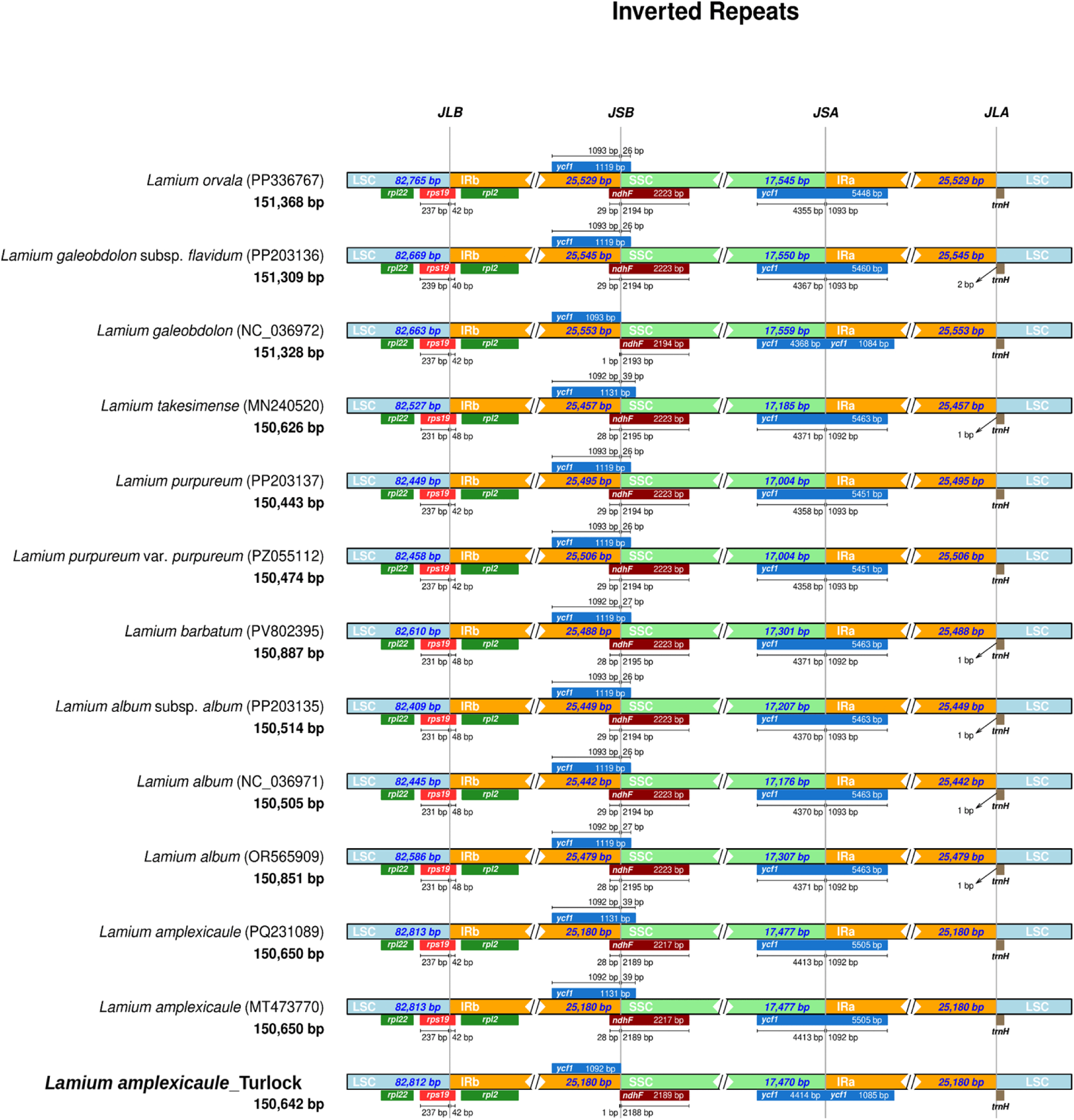
Comparison of inverted-repeat boundaries among 13 *Lamium* chloroplast genomes. JLB, LSC/IRb junction; JSB, IRb/SSC junction; JSA, SSC/IRa junction; JLA, IRa/LSC junction. Gene positions and region lengths are shown for each accession.

### 3.5 Sliding-window nucleotide diversity

The mean nucleotide diversity across the 13 *Lamium* plastomes was π = 0.01025 (Fig 5). The highest nucleotide-diversity value occurred in *ycf1* (π = 0.04314), followed by the *petA-psbJ* region (π = 0.04049).

**Fig 5.**
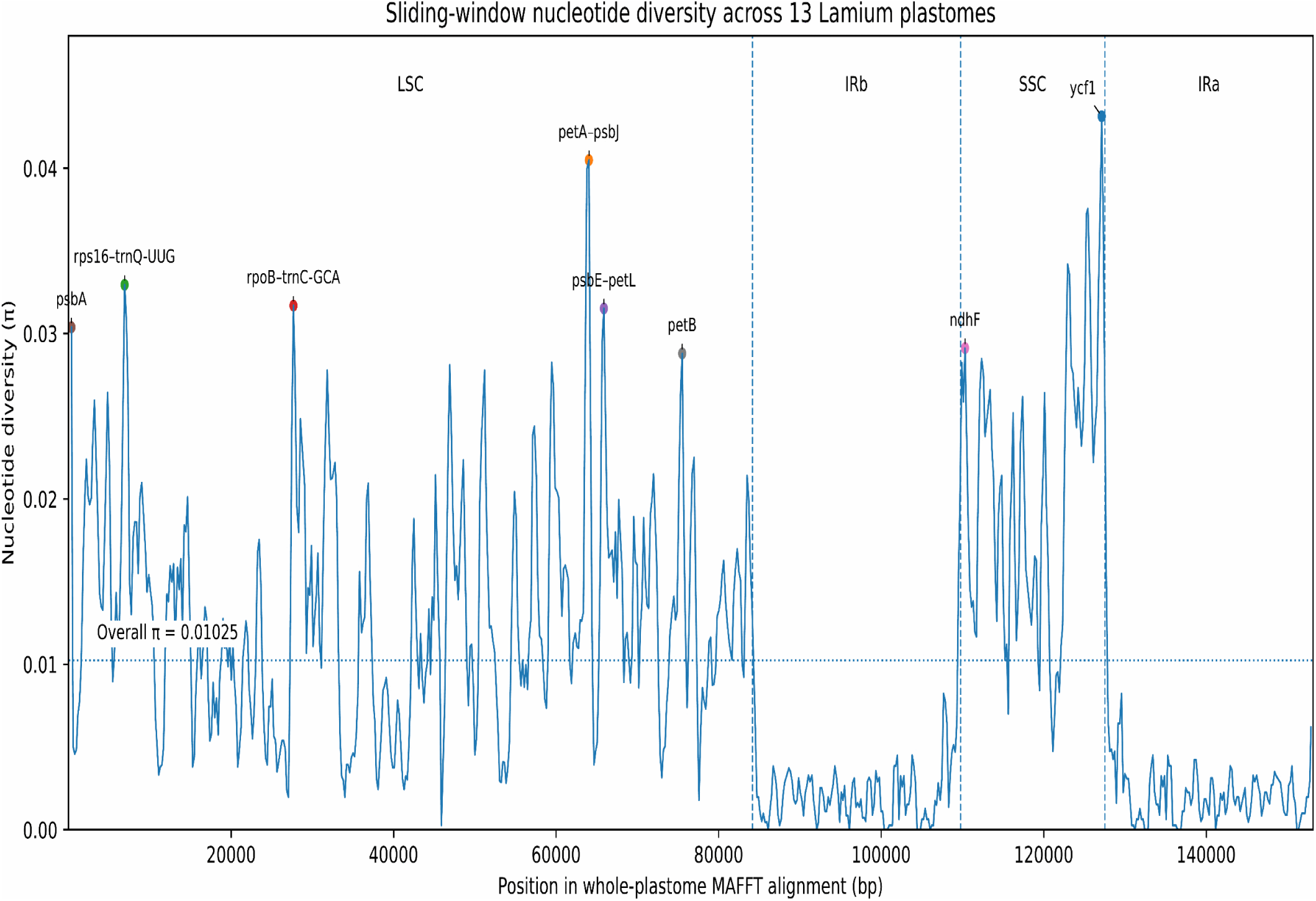
Sliding-window nucleotide diversity across 13 *Lamium* plastomes. Nucleotide diversity (π) was calculated using a 600-bp window and 200-bp step. Dashed vertical lines show the LSC, IRb, SSC, and IRa boundaries, and the horizontal dotted line shows the overall π value of 0.01025.

Other regions with high nucleotide diversity included *rps16-trnQ-UUG* (0.03295), *rpoB-trnC-GCA* (0.03169), *psbE-petL* (0.03152), *psbA* (0.03038), *ndhF* (0.02912), and *petB* (0.02880). The *ycf1* and *petA-psbJ* regions were selected for further marker analysis.

### 3.6 Repeat sequence variation

A total of 607 SSRs were identified across the 13 *Lamium* plastomes (Fig 6). Mononucleotide repeats were the most common, with 367 repeats (60.5%), followed by 84 dinucleotide, 78 tetranucleotide, 66 trinucleotide, eight hexanucleotide, and four pentanucleotide repeats. The number of SSRs ranged from 36 in *L. barbatum* to 56 in the *L. amplexicaule*_Turlock plastome. MT473770 and PQ231089 each contained 54 SSRs. Long dispersed repeats ranged from 20 to 24 per plastome after the main inverted repeats were excluded. All three *L. amplexicaule* plastomes contained 22 dispersed repeats. Tandem-repeat numbers ranged from 15 to 23, with 20 detected in each of the three *L. amplexicaule* plastomes. Most tandem repeats had periods between 11 and 30 bp.

**Fig 6.**
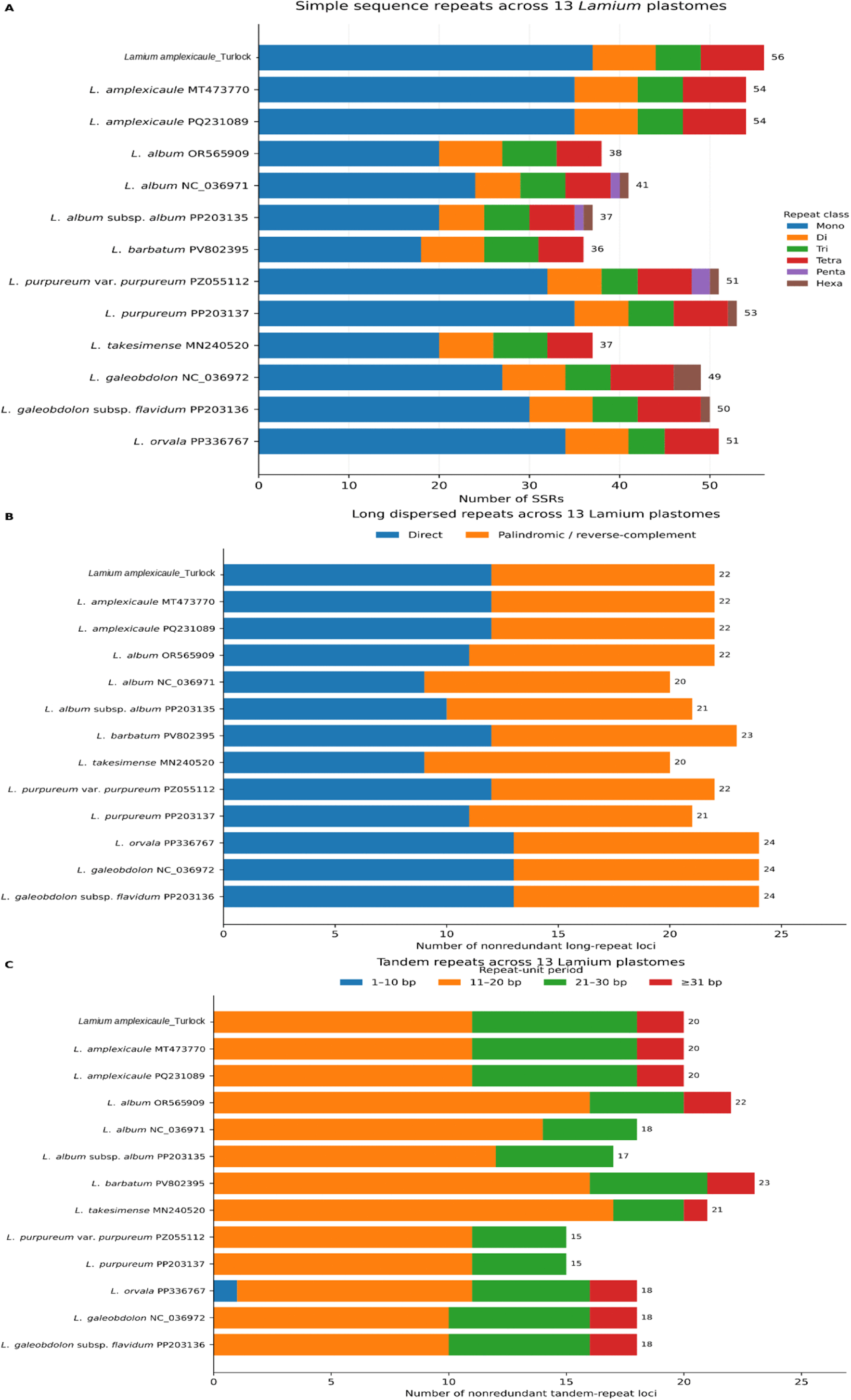
Repeat sequences across 13 *Lamium* plastomes. (A) Simple sequence repeats. (B) Long dispersed repeats. (C) Tandem repeats grouped by repeat-unit length. Values at the ends of the bars indicate total repeats or repeat loci.

### 3.7 Codon-usage patterns

Codon usage was similar among the 13 *Lamium* plastomes (Fig 7). Effective number of codons (ENC) values ranged from 50.24 to 50.47, with a mean of 50.41. Coding-sequence GC content ranged from 38.5% to 39.1%. Mean GC1, GC2, GC3, and GC3s values were 46.95%, 39.12%, 30.82%, and 27.78%, respectively. Thirty codons had mean RSCU values greater than 1.0, and 28 of these ended in A or U. UUA, which encodes leucine, had the highest mean RSCU value at 1.902. Overall, the preferred synonymous codons were mainly A- or U-ending codons.

**Fig 7.**
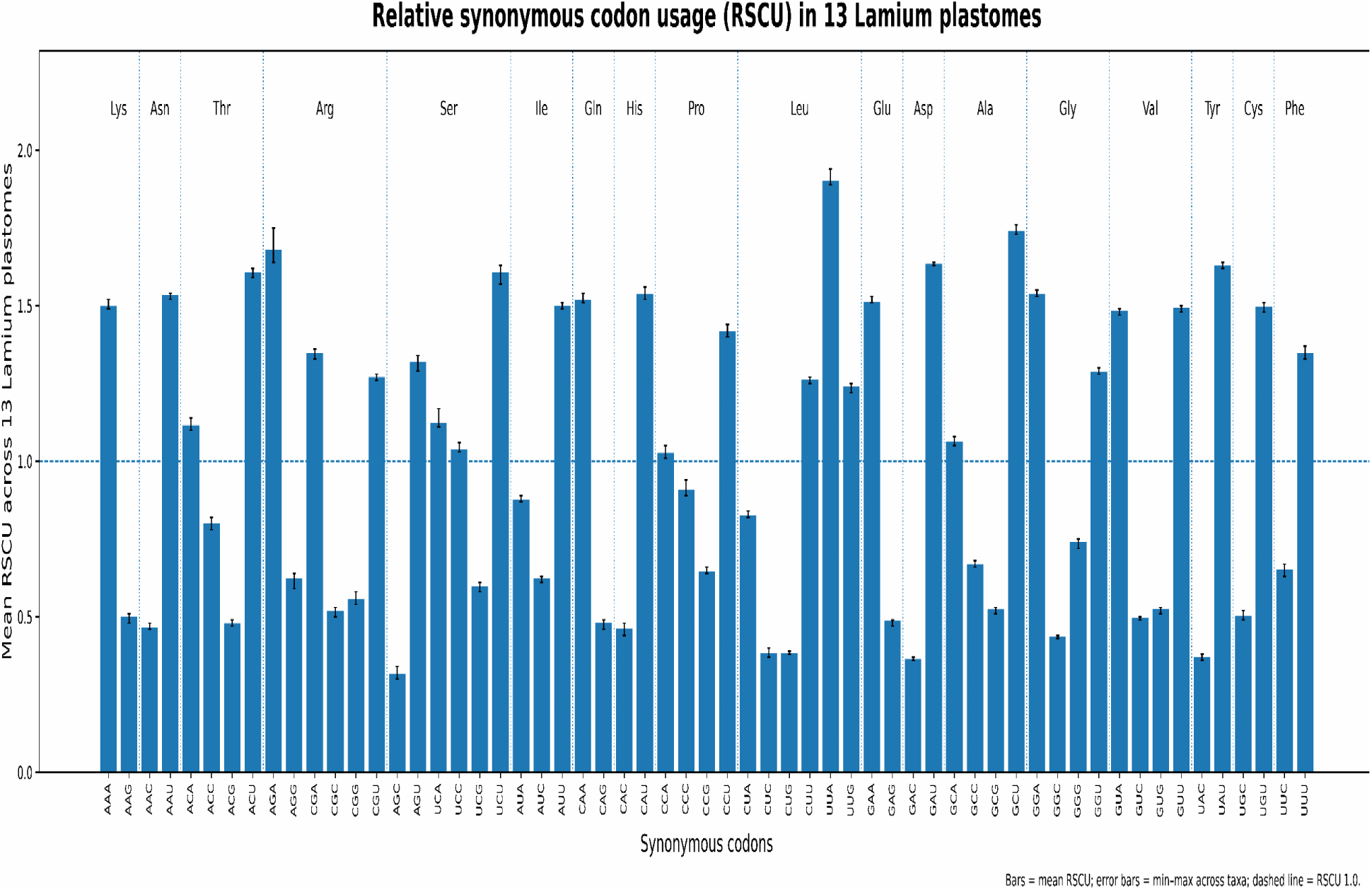
Relative synonymous codon usage across 13 *Lamium* plastomes. Bars show mean RSCU values from 78 shared protein-coding genes, and error bars show the minimum–maximum range among accessions. The dashed line indicates RSCU = 1.0.

### 3.8 SNP and indel variation

Across the 13-*Lamium* whole-plastome alignment, 4,004 complete-case SNP sites were identified. These included 3,888 biallelic and 116 multiallelic sites. DnaSP identified 726 indel events, with a mean inferred length of 11.03 bp. A total of 36 SNPs were found among the three *L. amplexicaule* plastomes. Fourteen SNPs occurred in introns, 12 were intergenic, and 10 occurred in protein-coding regions. Of the coding SNPs, five were synonymous and five were nonsynonymous. Coding SNPs occurred in *rpoA*, *ycf2*, *ndhF*, *ndhA*, and *ycf1*. One nonsynonymous *ycf2* substitution occurred in both IR copies. A cluster of 12 SNPs was also found in the *petD* intron.

DnaSP identified 16 indel events among the three *L. amplexicaule* plastomes. Mapping of the alignment gaps resolved these into 15 physical loci covering 24 gap positions. Twelve loci were intergenic and three were intronic, occurring in the *rps16*, *atpF*, and *pafI* introns. No indels were found within protein-coding genes, tRNAs, or rRNAs.

### 3.9 Selection pressure based on Ka/Ks

Ka/Ks values were calculated for 6,084 gene-by-pair comparisons across the 13 *Lamium* plastomes (Fig 8). Of these, 4,204 produced numeric ratios and 1,880 could not be estimated. Among the numeric comparisons, 4,169 had Ka/Ks values below 1 and 35 had values above 1. A total of 1,682 comparisons had Ka/Ks < 1 and Fisher *P* < 0.05. None of the 35 comparisons with Ka/Ks > 1 had a significant Fisher test. No significant evidence of positive selection was therefore detected.

**Fig 8.**
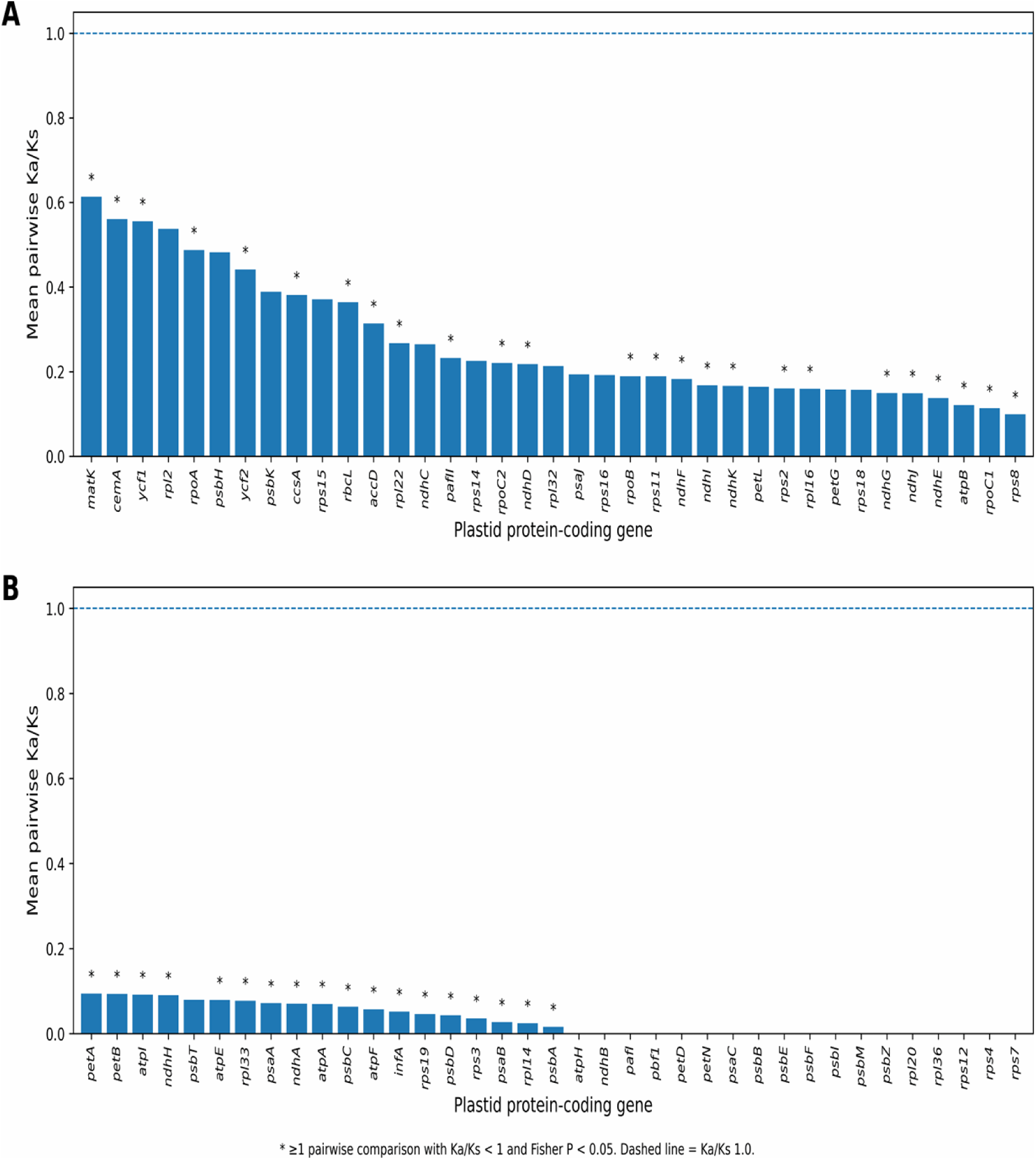
Mean pairwise Ka/Ks ratios for chloroplast protein-coding genes across 13 *Lamium* plastomes. Genes are divided between panels A and B for readability. Bars show mean pairwise MYN estimates, and the dashed line indicates Ka/Ks = 1.0. Asterisks indicate genes with at least one pairwise comparison showing Ka/Ks < 1 and Fisher P < 0.05. Genes without calculable ratios are not shown.

Numeric Ka/Ks estimates were obtained for at least one comparison in 74 of the 78 genes, and all 74 genes had mean Ka/Ks values below 1. The highest mean ratios were found in *matK* (0.614), *cemA* (0.561), *ycf1* (0.555), *rpl2* (0.537), *rpoA* (0.488), *psbH* (0.482), and *ycf2* (0.441). No calculable ratios were obtained for *psaI*, *psbJ*, *psbL*, or *rpl23*.

### 3.10 In silico barcode evaluation

Eight nucleotide-diversity hotspots were tested as 600-bp candidate marker regions across the 13 *Lamium* plastomes, representing seven species (S2_Table). The *ycf1* region contained 69 variable sites and 60 parsimony-informative sites and produced 11 haplotypes. The *petA-psbJ* region contained 58 variable sites and 53 parsimony-informative sites and produced 10 haplotypes (Table 2).

**Table 2.** In silico evaluation of candidate barcode regions in *Lamium*.

| Region | Variable sites | Parsimony-informative sites | Haplotypes | Interspecific identical haplotypes | Predicted amplicon | Barcode gap |
|---|---|---|---|---|---|---|
| *ycf1* | 69 | 60 | 11 | None | 556 bp | No |
| *petA-psbJ* | 58 | 53 | 10 | None | 511-561 bp | No |

No identical complete-case haplotypes were shared between different species for either *ycf1* or *petA-psbJ*. Both regions produced species-specific haplotypes for all seven sampled species. The other six candidate regions contained at least one identical haplotype shared between species. Neither *ycf1* nor *petA-psbJ* showed a positive classical barcode gap. Primer analysis identified one compatible forward and reverse binding site in all 13 plastomes for both regions. The *ycf1* primers produced a predicted 556-bp amplicon containing 64 variable sites, 55 parsimony-informative sites, and 11 haplotypes. The *petA-psbJ* primers produced predicted amplicons ranging from 511 to 561 bp, with 58 variable sites, 53 parsimony-informative sites, and 11 haplotypes. No exact interspecific sequence matches were found for either predicted amplicon. *ycf1* was retained as the primary candidate region and *petA-psbJ* as the secondary candidate.

## 4. Discussion

### 4.1 Plastome conservation and structural variation in *Lamium*

The chloroplast genome of the *L. amplexicaule*_Turlock accession was similar in size and structure to other published *Lamium* plastomes. The genome was 150,642 bp and had the typical quadripartite structure found in most land plants [1]. The previously reported *L. takesimense* plastome was 150,626 bp [6], while the *L. barbatum* plastome was 150,887 bp [7]. These results show that chloroplast genome size and general structure are highly conserved within *Lamium*. The standardized annotation also showed the same set of 113 distinct chloroplast genes in all 13 *Lamium* plastomes. Some earlier *Lamium* plastome studies reported different total gene numbers [6,7]. This may partly result from differences in how duplicated IR genes, partial genes, and tRNAs were annotated. Reannotating the plastomes using the same method reduced these differences and showed no gene presence-absence variation among the accessions examined here.

The synteny and IR-boundary analyses also showed that genome structure was mostly conserved. The main difference was the opposite SSC orientation in the *L. amplexicaule*_Turlock plastome. Different SSC orientations have been reported in chloroplast genomes and can result from recombination between the inverted repeats [36,37]. Because no other major rearrangements were detected, the SSC orientation in the *L. amplexicaule*_Turlock accession is more likely an alternative structural form of the plastome than a fixed inversion unique to this accession.

### 4.2 Phylogenomic relationships and locus-level concordance

The phylogenomic analysis strongly supported the sampled *Lamium* accessions as a clade. The three *L. amplexicaule* plastomes were also grouped together, with the *L. amplexicaule*_Turlock accession sister to MT473770 and PQ231089. This agrees with the broader Lamiaceae plastome phylogeny reported by Zhao et al. [2] and supports the placement of the *L. amplexicaule*_Turlock accession within *L. amplexicaule*. Several relationships within *Lamium* were also similar to earlier plastome studies. *Lamium takesimense* has previously been placed near *L. album* [6], and a more recent study placed *L. barbatum* in the same general group as *L. takesimense*, *L. album*, *L. amplexicaule*, and *L. galeobdolon* [7]. In the present analysis, *L. barbatum* and *L. takesimense* were recovered as sister taxa, with the *L. album* accessions nearby. However, the chloroplast tree should not be considered a complete species phylogeny for *Lamium*. Earlier work using both chloroplast and nuclear markers found disagreement between the two genomes and suggested that hybridization and polyploidy have played an important role in the evolution of the genus [4]. Similar disagreement between plastid and nuclear phylogenies has also been reported elsewhere in Lamioideae [38]. Nuclear genome data will therefore be needed to determine whether the relationships found here are also supported by independently inherited markers.

The gene concordance factors also showed that support was not the same across individual genes. Several branches had very high bootstrap support but lower gCF values. These measures are not expected to give identical results because bootstrap support measures support in the concatenated dataset, while gCF measures how many individual gene trees contain the same branch [13,39]. The removal of *clpP1* did not change any internal branches in the tree and caused only small changes in gCF values. This indicates that the *clpP1* issue did not have an important effect on the main phylogenomic result.

### 4.3 Sequence divergence and mutational hotspots

Although the *Lamium* plastomes were highly conserved in overall structure, nucleotide variation was not evenly distributed across the genome. Several of the most variable regions in this study have also been reported as variable chloroplast regions in other plants. These include *petA-psbJ*, *rps16-trnQ*, *psbE-petL*, *ndhF*, and *ycf1* [9,10].

The highest nucleotide-diversity value occurred in *ycf1*, followed by the *petA-psbJ* region. Previous studies have also identified *ycf1* as a highly variable plastid region with useful species-level discrimination [11]. The occurrence of several previously reported variable regions among the highest peaks in *Lamium* suggests that these regions may also be useful for studying closely related taxa within this genus.

Variation was also present among the three *L. amplexicaule* accessions. A total of 36 SNPs and 16 inferred indel events were found even though the three plastomes were very similar in genome structure and gene content. These differences show that sequence-level variation is present within the sampled *L. amplexicaule* plastomes. However, only three accessions were available, so the observed differences cannot yet be used to determine geographic or population-level patterns. More samples from different populations would be needed for that type of comparison.

### 4.4 Repeat composition, codon usage, and selective constraint

Repeat sequences showed more variation among the *Lamium* plastomes than overall genome structure. Mononucleotide SSRs were the most common repeat type, which is consistent with chloroplast SSR patterns reported in many plants [8]. The total number of SSRs differed among accessions, including among the three *L. amplexicaule* plastomes. In comparison, the three *L. amplexicaule* accessions had the same numbers of long dispersed and tandem repeats. Chloroplast repeats may therefore provide useful markers for future studies.

Codon usage was much more similar across the genus. Most preferred synonymous codons ended in A or U, and GC3 and GC3s values were low. A preference for A/T-ending codons has also been reported in chloroplast genomes from other plant groups [40,41]. The codon-usage pattern found in *Lamium* therefore appears to follow the general pattern reported for plastid genomes rather than being specific to the genus.

The Ka/Ks results showed that most chloroplast protein-coding genes were under purifying selection. Most calculable comparisons had Ka/Ks values below 1, and all statistically significant comparisons were in the purifying-selection direction. A small number of comparisons had Ka/Ks values above 1, but none were statistically significant. These values were therefore not considered evidence of positive selection. Ka/Ks values can also be unstable when closely related sequences contain only a small number of substitutions [42], so the significant tests and overall pattern are more informative than isolated ratios above 1.

### 4.5 Candidate diagnostic regions and future directions

The two strongest candidate regions in the in-silico analysis were *ycf1* and *petA-psbJ*. Both regions have previously been reported as variable chloroplast loci [9–11]. In the present dataset, both regions distinguished the seven sampled *Lamium* species without exact sequence sharing between species. The predicted *ycf1* amplicon was also the same length across all 13 plastomes. However, neither region produced a positive classical barcode gap. The ability to distinguish the accessions included in this study does not mean that the same regions will distinguish all *Lamium* species or populations. This is especially important because previous work has shown complex relationships within *Lamium* involving polyploidy and hybridization [4]. The primers and candidate regions identified here should therefore be tested using more species, more accessions per species, and experimental PCR and sequencing. Additional sampling would also be useful within *L. amplexicaule*. The species has both cleistogamous and chasmogamous flowers, and its reproductive strategy can respond to environmental conditions [5]. Sampling populations from different geographic regions could help determine whether the sequence differences observed among the three available plastomes are associated with population-level variation. Nuclear sequence data would also allow comparison of the chloroplast phylogeny with an independently inherited genome, which is important because plastid-nuclear disagreement has previously been reported in *Lamium* and related groups [4,38].

The chloroplast genomes examined here were highly conserved in structure and gene content, but variation was present in nucleotide sequence and repeat composition. The *L. amplexicaule*_Turlock accession was placed with the other *L. amplexicaule* plastomes, and this placement was unchanged after removing *clpP1*. The variable regions identified here, especially *ycf1* and *petA-psbJ*, provide useful candidates for future work with larger *Lamium* datasets.

## 5. Conclusion

In this study, we characterized the complete chloroplast genome of *Lamium amplexicaule* collected in Turlock, California, and compared it with other available *Lamium* plastomes. The *Lamium amplexicaule*_Turlock plastome was 150,642 bp and showed the typical quadripartite structure. Gene content and overall genome organization were highly conserved across the 13 *Lamium* plastomes. Phylogenomic analysis placed the three sampled *L. amplexicaule* accessions together in a strongly supported clade, and removal of *clpP1* did not change the tree topology. The three *L. amplexicaule* plastomes also showed sequence-level differences, including 36 SNPs and 16 inferred indel events.

Most protein-coding genes showed evidence of purifying selection, with no significant evidence of positive selection. The *ycf1* and *petA-psbJ* regions showed the highest nucleotide diversity and were the strongest candidate markers in the in-silico analysis, although additional sampling and experimental testing are still needed. The results show that *Lamium* chloroplast genomes are highly conserved in structure and gene content but contain sequence variation that may be useful for future phylogenetic, taxonomic, and population studies.

## Acknowledgment

This study was conducted as part of the Course-based Undergraduate Research Experience (CURE) Project through BIOL 4980, offered during the Fall 2026 semester. We thank the students and laboratory support staff who contributed to the successful completion of this research.

## Supporting information

S1_Data. Supporting data underlying the comparative plastome analyses. Includes taxon information, candidate-marker comparisons, repeat analyses, codon-usage data, Ka/Ks summaries, focal SNP and indel data, and in silico primer analyses.

S1_Table. Taxa and chloroplast genome accessions used in this study.

S2_Table. In silico evaluation of eight candidate plastid marker regions in Lamium.

S1_Code. Validated reconstruction of the custom Python workflow used for in silico Lamium barcode evaluation and primer screening.

